# Visuomotor flexibility is embedded in the topography of frontal cortex

**DOI:** 10.64898/2026.05.25.727601

**Authors:** Nicholas M. Dotson, John H. Reynolds

## Abstract

Visual selection and saccade targeting are not obligatorily coupled, allowing a stimulus at one location to guide an eye movement toward another. This is mediated by frontal oculomotor cortex, where maps of visual and motor space converge, yet whether the topographic relationship between these maps itself contributes to this flexibility is unknown. Using Neuropixels recordings across horizontal segments of marmoset cortex, we find that individually visual and saccade vector angles change smoothly, with occasional abrupt jumps. Rather than a one-to-one visual-motor mapping, local cortical patches contain a combination of aligned and misaligned vector angles. Mosaic map models reproduce the empirical distributions of local angle differences for visual and motor maps, revealing that the two maps have constrained and distinct spatial scales. When combined, these mosaic maps generate a moiré pattern that quantitatively matches the observed visual-motor angular differences, providing a substrate for flexible transformations between visual inputs and saccade outputs.

---

Flexible gaze control requires that visual signals be selectively routed to different motor outputs depending on behavioral context. How this flexibility is implemented in the neural circuits that link vision to action remains poorly understood. The frontal eye fields (FEF) and adjacent premotor eye fields (collectively FEF+) are key visuomotor hubs for controlling gaze and directing attention (Robinson & Fuchs, 1969; Bruce et al., 1985; Hanes & Schall, 1996; Sommer & Tehovnik, 1997; Moore et al., 2003; Schiller & Tehovnik, 2005; Lowe et al., 2022). Electrical microstimulation at nearby sites across the cortex in FEF produces smooth changes in saccade vector angles, interrupted by abrupt jumps and repetitions (Bruce et al., 1985). The angular position of visual receptive fields (RFs) demonstrates a similar pattern (Suzuki & Azuma, 1983; Suzuki, 1985). These observations are not compatible with a simple retinotopic map. Rather, they suggest a more complex topography. We posited that the vector angles of saccade motor fields (MFs) and visual RFs are organized in a mosaic-like topography analogous to the direction-selective maps observed in ferret V1 (Weliky et al., 1996), which would produce locally smooth changes in vector angles punctuated by abrupt discontinuities across the cortex (Fig. 1A and B).

**Figure 1.**
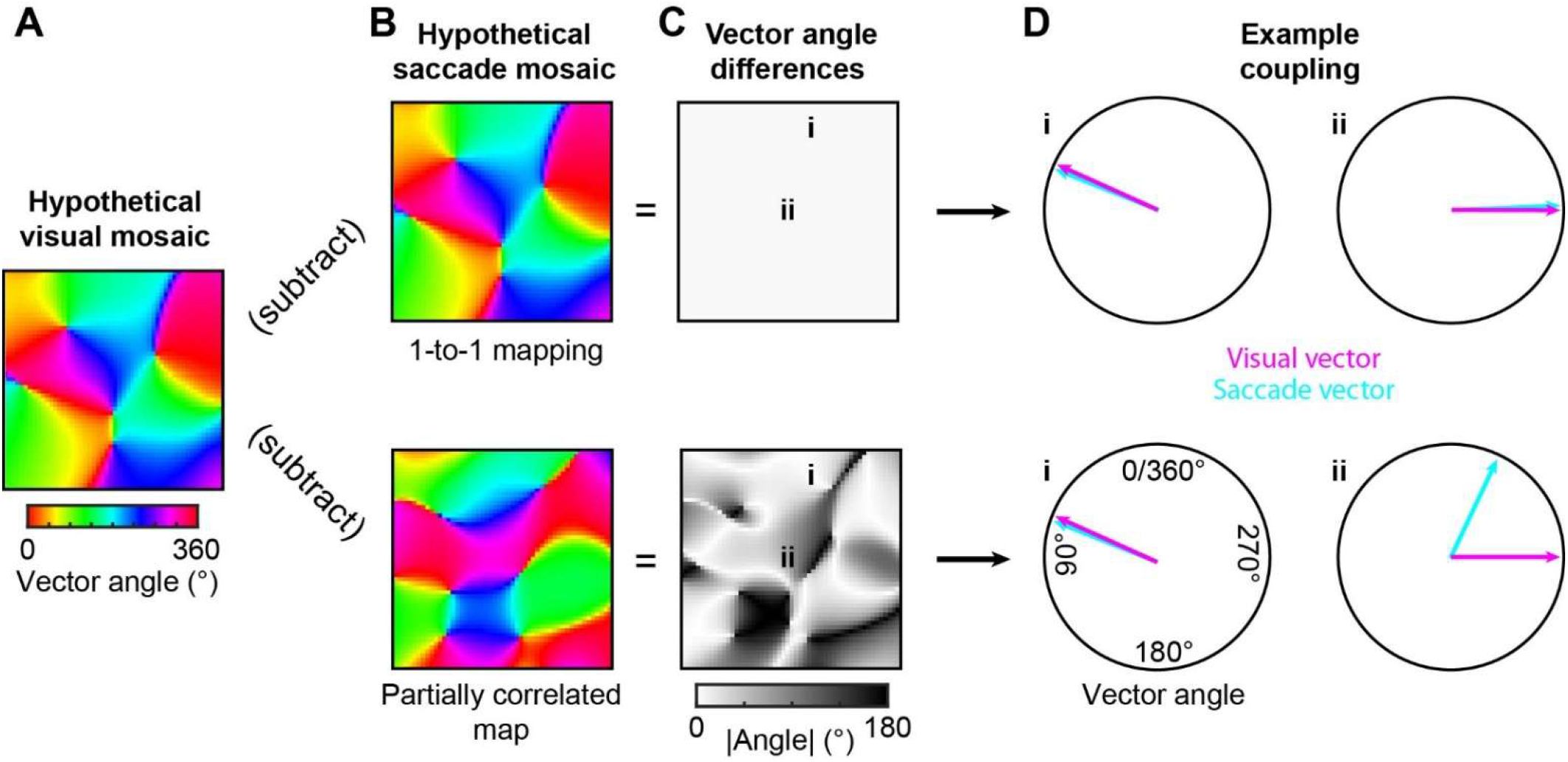
Moiré visuomotor topography as a mechanism for flexible visual-motor coupling. **A, B**. Hypothetical mosaic map topography for visual RF and saccade MF vector angles. The vector angle is the angle of the RF or MF in a retinotopic reference frame. Each pixel on the maps corresponds to a physical region on the cortex. Color indicates the vector angle (visual or saccade) at that position. **C**. Difference between visual and saccade mosaic maps. If maps are identical (top) vectors are always aligned, whereas partially correlated maps enable multiple vector combinations. **D**. For a 1-to-1 coupling scheme (top), vectors are always aligned (i, ii). When mosaics are only partially correlated (bottom), vectors can both align (i) and diverge (ii).

Visual and saccade target signals are generally thought to align, however, visual selection and saccade target selection are not obligatorily linked (Hallett, 1978; Thompson et al., 1997; Murthy et al., 2001). This dissociation implies that the neural substrate supporting visuomotor transformations should permit flexible relationships between visual and motor representations. A strictly one-to-one topographic coupling between RF and MF vectors would constrain such flexibility (Fig. 1 top row). We therefore hypothesized that visuomotor topography is only partially coupled in FEF+. In this framework, non-identical but overlapping mosaic maps could support flexible coupling between visual locations and saccade targets across a broad range of angular combinations, while maintaining regions of alignment (Fig. 1 bottom row). We further propose that this flexibility arises from a constrained mismatch in spatial scale between the visual and motor mosaics, such that their interaction generates a structured moiré interference pattern of RF–MF pairings. The spatial pattern of alignment and misalignment should be observable as a moiré interference pattern in the RF-MF angle differences across the cortical surface (Fig. 1C), and should, in principle, support the range of vector transformations required for tasks such as anti-saccades.

## Results

To map visuomotor topography, we leveraged two key advantages of the marmoset. First, FEF+ lies on the surface of the lissencephalic (smooth) marmoset cortex, unlike the macaque where it is buried in the arcuate sulcus, enabling direct access for high-density recordings across the cortical surface (Collins et al., 2005; Selvanayagam et al., 2019; Kaas, 2020; Feizpour et al., 2021; Dotson et al., 2024). Second, the compact representation of visuomotor space in marmoset FEF+ produces large changes in vector angles over short horizontal distances, making it possible to characterize topographic structure with laminar probes. We mapped visual RFs and saccade MFs of individual neurons using Neuropixels probes inserted at an average angle of 18.2° ± 8.3° (mean ± SD) to record across horizontal segments of cortex (Fig. 2A, B). Oblique rather than perpendicular penetrations were used because, while perpendicular penetrations sample neurons within the same functional column across layers, angled penetrations traverse the cortical surface and thereby track changes in tuning across space. This approach previously revealed mosaic or pinwheel-like organization of direction preferences in macaque MT (Albright et al., 1984).

**Figure 2.**
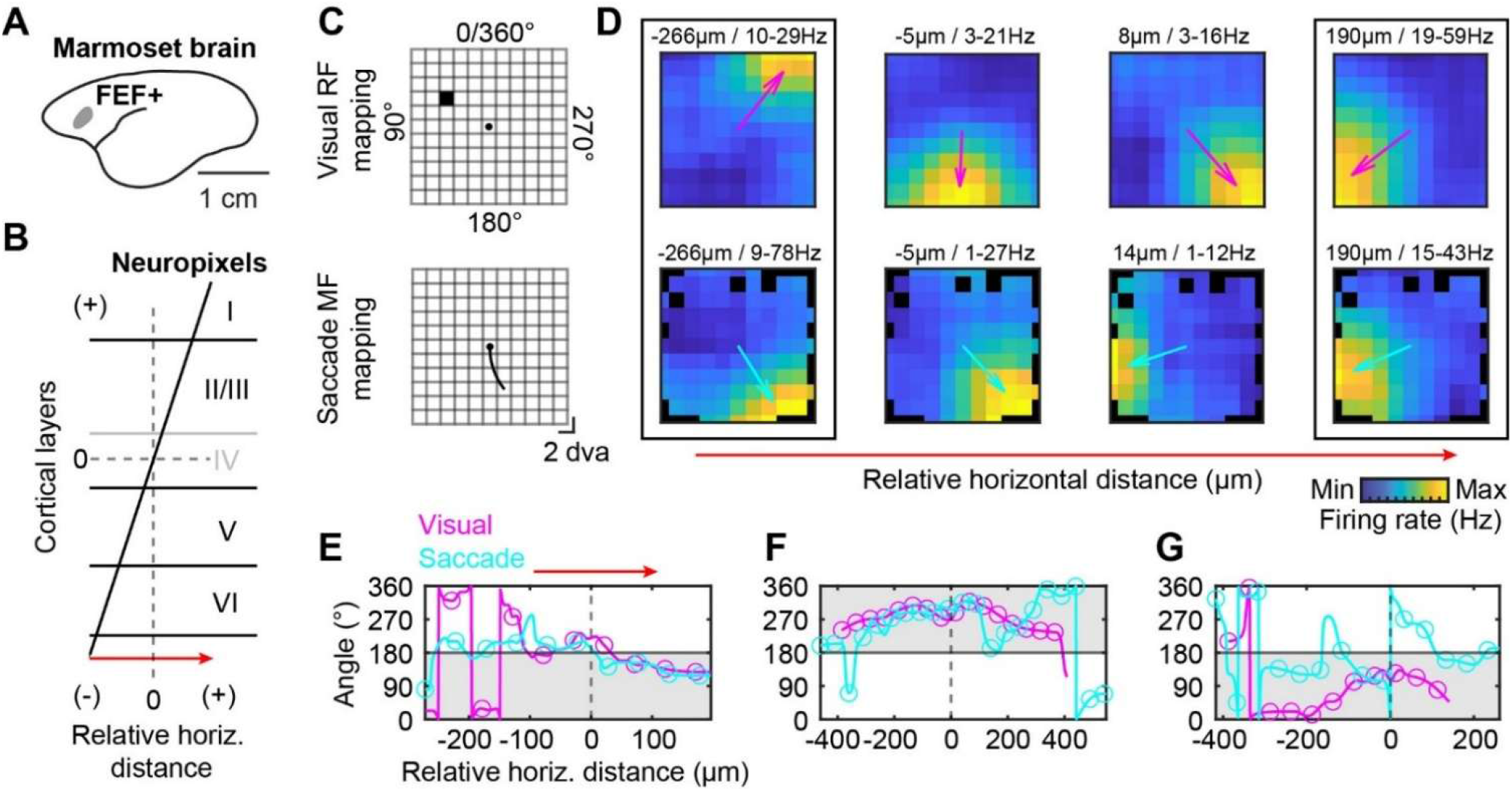
Visual RF and saccade MF vector angles are semi-independent and change in a manner consistent with a mosaic topography. **A**. Illustration of the approximate location of FEF+ on the marmoset brain. **B**. Illustration of a Neuropixels probe passing through the cortex at an angle. Cortical layers are labeled. Layer IV is in light gray to note that its presence varies across the frontal cortex. **C**. Illustration of the RF and MF mapping procedure. **D**. Examples of RFs (top) and MFs (bottom) recorded simultaneously (same data as E). The far left example shows a neuron with different RF and MF vectors. The middle two columns show examples of nearby neurons with differing RFs and MFs. The far right example shows a neuron with highly similar RF and MF vectors. The relative horizontal location and the minimum and maximum firing rates are indicated at the top of each example. Black rectangles indicate that the RF and MF are from the same neuron. For visualization purposes data were smoothed by 2 bins. **E-G**. Examples of the RF and MF vector angles across short horizontal segments of FEF+. The data shows the interpolated values across the cortex. Circles indicate 50 micron steps, which are used for further analyses. Shaded regions indicate the contralateral hemifield.

RF mapping was performed by flashing a small stimulus (2dva) at randomized locations covering 22 by 22 degrees of visual angle (dva) (Fig. 2C). The mapping task did not require maintaining constant fixation, enabling us to utilize saccades made during the task to identify MFs. However, only visual stimuli flashed during periods of fixation were used, and the position of the stimuli were corrected for eye position. Similar “free-viewing” tasks have been successfully utilized to quantify visual RFs in visual cortical areas (Yates et al., 2023). We recorded data from a total of 39 sites in two marmosets, comprising 4,221 isolated units (Supp. Fig. 1A). Of these, 2,783 and 3,011 units were sufficiently active during visual stimulation and saccade production, respectively, and were included in the analyses. We used Spatial Information Analysis (Skaggs et al., 1996; Dotson & Yartsev, 2021) to identify neurons with RFs (22%), MFs (31%), or both (10%). We found that only sites within the putative FEF+ (18/39) region had a high incidence of RFs and MFs (Supp. Fig. 1A; Selvanayagam et al., 2019). Subsequently, all further analyses considered only these sites.

### Maps are partially coupled

We found a high incidence of RFs and MFs across all laminar compartments (∼10 to 40%; Supp. Fig. 1B, C), consistent with visuomotor integration across cortical layers. Each RF and MF was assigned a vector, and the angle of each vector was used for subsequent analyses (Fig. 2D). Individual neurons could have similar RF and MF angles or substantially different ones, and RF and MF locations could differ even for the same neuron (Fig. 2D; Supp. Fig. 1E, F). In agreement with prior studies (Crapse & Sommer, 2009), RFs and MFs were present for both the contralateral and ipsilateral hemifields (summary is shown Supp. Fig. 1D and E split by animal). In some cases, neurons have similar RFs and MFs, while in others the vector angles differ substantially. RF and MF locations can even differ for the same neuron (summary is shown in Supp. Fig. 1E and F split by animal). To visualize the changes in angles across the cortex and to format the data in a manner that can be compared across penetrations and modeled, RF and MF angles were locally averaged (± 25µm), and then interpolated and resampled at 50µm intervals (Fig. 2E-G; Supp. Fig. 1H). RF and MF angles change smoothly with occasional jumps and drift with respect to one another across the cortical surface (Fig. 2E-G), indicating partial rather than rigid coupling and raising the question of whether this drift reflects an underlying difference in the spatial organization of the two maps. These observations motivate a quantitative analysis of the spatial scales of the visual and motor mosaics, to test whether a specific mismatch in scale can account for the observed RF-MF angle relationships and thus implement the proposed moiré visuomotor topography.

### Maps have a mosaic topography

To test our hypothesis that RF and MF vector angles are organized into mosaic maps, we compared observed differences in neighboring vector angles with those simulated using mathematical mosaic map models. Our aim was to determine whether the spatial structure of each map can be captured by a narrow range of spatial scales, as would be expected if the visual and motor mosaics form the building blocks of a structured moiré topography. We started with a model that generates mosaic maps by summing plane waves whose orientations and spatial frequencies are determined by an annulus in 2D Fourier space (Fig. 3A, B; Niebur & Wörgötter, 1994; Wolf & Geisel, 1998). The annulus is defined by a center spatial frequency (SF) and a bandwidth (BW), which together control the range of spatial frequencies. This type of annulus mosaic (AM) model produces biologically realistic mosaic maps. We find that the rate-of-change map is highly similar to that observed for direction-selectivity maps in ferret V1, accounting for the abrupt jumps in tuning angle observed experimentally (Supp. Fig. 2; Weliky et al., 1996).

**Figure 3.**
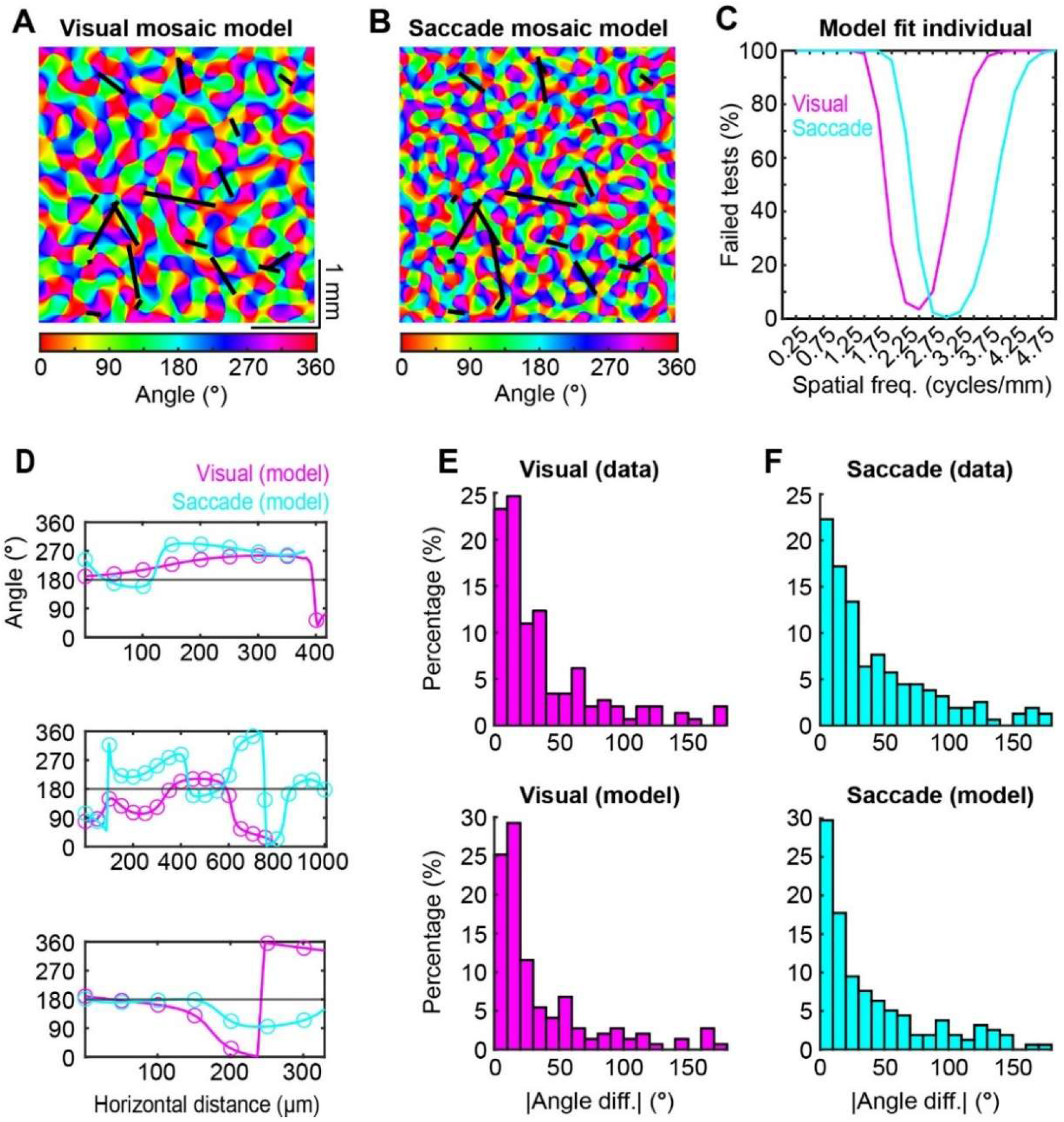
Visual and saccade mosaic maps match data, but with different preferred spatial frequencies. **A, B**. Examples of randomly generated visual (A) and saccade (B) mosaic maps. Black lines indicate the sampled locations (n = 18). Line lengths match the horizontal distances between the first and last tuned neuron for each probe penetration. In some instances this distance was shorter for either the visual or saccade data. **C**. Results from testing the distribution of differences from the model to the real data at a range of spatial frequencies. Only models within a narrow but offset range of spatial frequencies fit the data. **D**. Examples of the visual and saccade angles from the models (A, B). Similar to the real data, there are smooth changes and abrupt jumps. Moreover, like the real data visual and saccade angles tend to change semi-independently. **E, F**. The distribution of differences for the real visual and saccade data (top) can be replicated by a mosaic map model (bottom). Model data is from a single simulation (same data as A, B, and D).

The AM model was simulated over a 4mm x 4mm area. To mimic how we sampled the cortex, the AM model was sampled at 50µm steps along randomly placed lines (18 lines) whose lengths matched the relative horizontal distances from the real data (Fig. 3A, B). The distribution of angular differences between neighboring samples along each randomly placed line was subsequently used to fit the models. Visual and saccade maps were best fit by distinct spatial frequencies with correspondence between model and data occurring only within a narrow SF band (Fig. 3C-F). Critically, the preferred SFs differed between maps, indicating that visual and motor mosaics are organized at distinct spatial scales. Figure 3A and 3B show examples of mosaic maps that closely fit the visual (SF = 2.25 cycles/mm) and saccade (SF = 3 cycles/mm) data. We determined which range of SFs best matched the empirical data by comparing the distribution of angle differences from the real data to those obtained from simulations (1,000 per SF combination) using a two-sample Kolmogorov-Smirnov test. SF values were considered consistent with the data if the simulated and empirical distributions did not differ significantly (“failed test”; p ≥.05). The visual and saccade data were fit separately. Bandwidth had little effect on results (Supp. Fig. 3); we therefore fixed it at 0.3 for all subsequent analyses. We found a strong correspondence between the real data and the model data for only a small range of spatial frequencies (Fig. 3C). Interestingly, the preferred SFs differed between the visual and saccade maps (Fig. 3C). Figure 3D shows examples of angles from the single lines on the mosaic maps in Figure 3A and B, which closely resemble the patterns observed in the real data (Fig. 2E-G). The suitability of the mosaic map model is further illustrated by the high degree of similarity between the distributions of angle differences derived from the example visual and saccade maps and the real difference distributions (Fig. 3E, F).

To test whether these findings depend on the specific architecture of the AM model, we repeated the analysis using an alternative noise mosaic (NM) model that generates irregular, puzzle-piece-like tuning patches (Supp. Fig. 4A, B). The NM model generated mosaic maps by convolving a complex Gaussian white-noise field with a 2D Gaussian kernel, with spatial scale controlled by the kernel width. Preferred spatial scales again occurred within narrow bands and were offset between visual and motor maps, with a lower preferred spatial scale for visual angles (Supp. Fig. 4C). The agreement between architecturally distinct models indicates that the mosaic organization of RF and MF angles reflects a general property of the data rather than an artifact of a particular model choice. The consistent finding that visual and motor mosaics prefer distinct spatial scales raises the question of whether this difference can account for the systematic drift between RF and MF angles.

**Figure 4.**
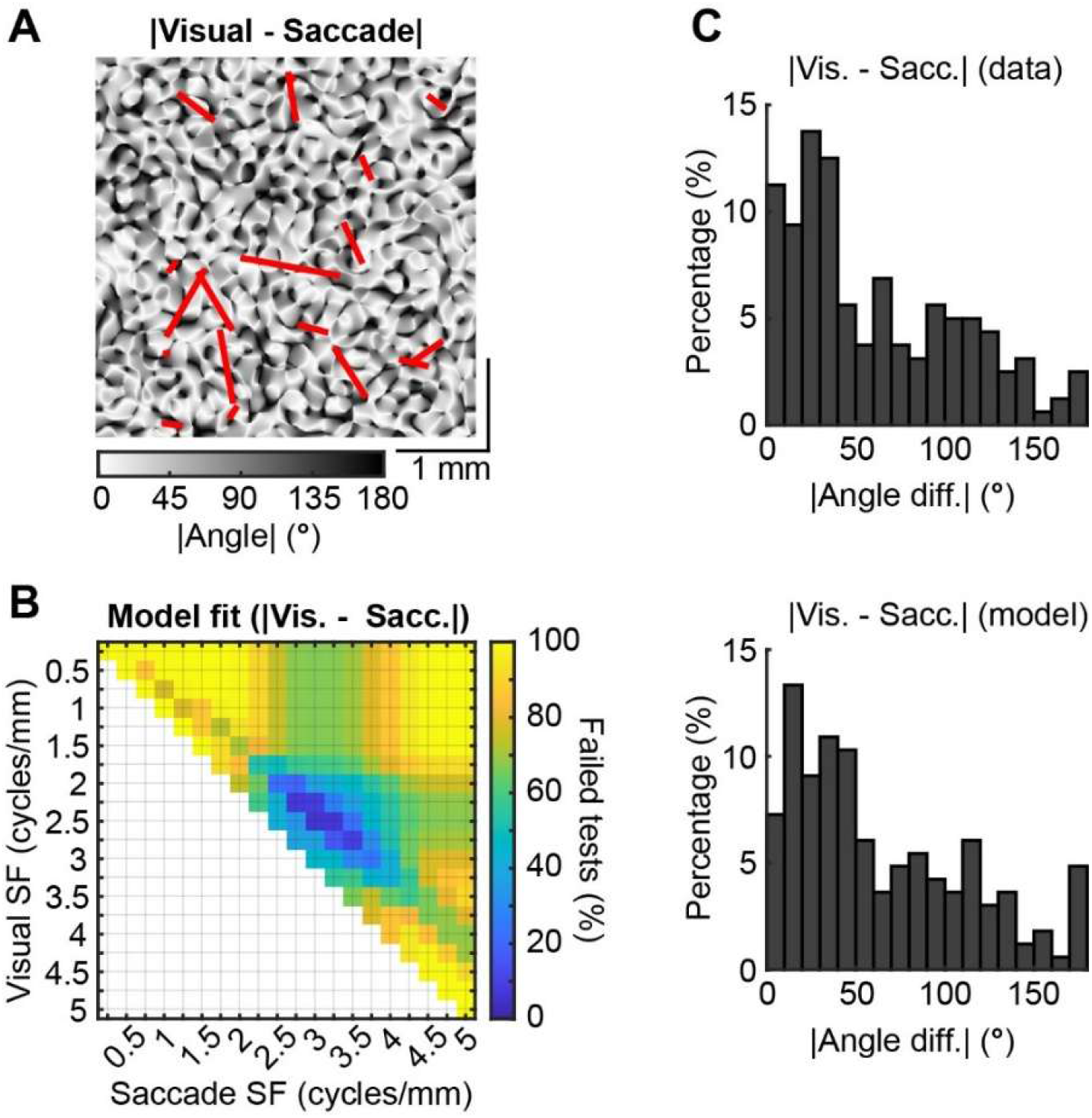
Moiré visuomotor topography accounts for the distribution of visual-motor angular differences. **A**. Example interference pattern generated by subtracting simulated visual and saccade mosaic maps with different SFs (Fig. 3A and B). The resulting difference map exhibits a spatially structured moiré pattern characterized by alternating regions of alignment and misalignment. Red lines indicate the sampled locations. The line lengths correspond to the line lengths of the visual data. **B**. Parameter sweep across visual and saccade SF combinations combined with the results from the individual model fits. Only a narrow band of SF combinations satisfied all three criteria simultaneously. Maps generated with identical SFs (diagonal) are necessarily identical given the shared phase initialization, producing zero angular differences by construction. **C**. Distribution of angular differences from the empirical data (top) compared with the distribution generated by the example simulation (bottom).

### Induced saccades have a mosaic topography

To independently test the mosaic organization of saccade motor vectors, we used electrical microstimulation along obliquely inserted laminar probes to induce saccades. Induced saccade angles changed smoothly across the cortex with occasional abrupt jumps (Supp. Fig. 5), consistent with the mosaic organization observed for MF angles and with prior microstimulation studies (Bruce et al., 1985). Induced saccade angles were consistent with mosaic topography (AM model) but at a lower preferred SF (∼1.5 cycles/mm) than unit-recorded MF angles (∼3 cycles/mm). This difference is expected: microstimulation at 25 µA engages neurons over a larger cortical radius than the ±25 µm averaging window used for MF angles, effectively blurring the spatial scale of the measurement (Stoney et al., 1968; Tehovnik, 1996). Rerunning the mosaic analysis for MF angles using a 100 µm averaging window reduced the preferred SF to the range observed with microstimulation (Supp. Fig. 6), confirming that the discrepancy reflects the spatial footprint of each measurement rather than a genuine difference in organization. The agreement between unit recording and microstimulation analyses across matched spatial scales provides methodologically independent support for the mosaic organization of saccade motor vectors.

### Moiré visuomotor topography

Could the difference in preferred spatial frequencies explain the apparent drift between the visual and saccade angles (Fig. 2E-G)? We therefore asked whether a moiré interference pattern, which arises from the superposition of two systematically offset lattice patterns, accounts for the observed differences between visual and saccade angles. In particular, we tested whether only a constrained band of spatial-frequency mismatches and coupling strengths between the visual and motor mosaics can reproduce the empirical RF-MF angle distribution, as predicted if this moiré structure embeds the available repertoire of visuomotor pairings. Two random mosaics, even with the same SF would produce entirely random differences in angles, not the semi-coupled patterns we observe. To isolate the contribution of spatial scale mismatch, we therefore generated visual and saccade maps that shared underlying structure but differed in SF. This was achieved by initializing both maps with the same random Fourier coefficients, producing partial overlap of their constituent plane waves due to the overlap between their annular frequency bands. Subtracting the visual and saccade mosaic maps (Figure 3A and B) reveals an undulating moiré interference pattern of alternating high- and low-coherence regions (Fig. 4A), consistent with the semi-independent relationship observed between visual and saccade angles.

To determine which combinations of visual and saccade SFs best accounted for the empirical RF-MF angular differences, we applied the same fitting procedure used for the individual maps. Each sampled RF-MF angular difference was used directly to construct the empirical distribution for model comparison. Systematic exploration of visual and saccade SF combinations revealed that only a restricted subset of parameter pairs reproduced the empirical distribution of angular differences (Fig. 4B). Figure 4B summarizes the intersection of three constraints: (1) parameter ranges that fit the visual map alone, (2) those that fit the saccade map alone, and (3) those that reproduced the RF-MF difference distribution. Only a narrow band of SF combinations satisfied all three criteria simultaneously. The angular difference distribution generated by the example simulation closely matches the empirical distribution (Fig. 4C). Together, these results show that a modest spatial frequency mismatch between partially correlated mosaics accounts for the systematic drift between visual and saccade angles. The narrowness of the parameter band satisfying all three constraints simultaneously argues that this correspondence reflects genuine topographic structure rather than a modeling artifact.

To test whether these conclusions depend on the specific AM model architecture, we repeated the analysis using the NM model. Unlike the AM model, the NM model requires spatial scale offset and correlation strength to be specified independently, allowing us to isolate the contribution of each property. Maps with distinct spatial scales and intermediate partial correlation (ρ ≈ 0.6) were required to reproduce the empirical RF-MF difference distribution, with fully independent and fully correlated maps both failing to do so (Supp. Fig. 4D, E). The convergence across architecturally distinct model classes indicates that these two properties reflect general requirements of the data rather than artifacts of any particular model choice. Together, the AM and NM model analyses establish that moiré visuomotor topography arises from a specific and constrained combination of distinct spatial scales and intermediate coupling, properties that are robust to the choice of mosaic model and therefore likely reflect the underlying organization of FEF+

## Discussion

Flexible visuomotor behavior requires that visual selection and saccade targeting be related but not obligatorily coupled (Hallett, 1978; Thompson et al., 1997; Murthy et al., 2001). Our findings show that this functional flexibility is embedded in the visuomotor topography itself: local patches of cortex are endowed with different combinations of visual and saccade vector angles, including both aligned and divergent pairings. This architecture is further supported by the difference in spatial scale between the two maps. Because the visual mosaic operates at a lower spatial frequency than the saccade mosaic, visual RF patches are spatially broader, meaning any given visual location is represented across a wider extent of cortex and therefore overlaps with a larger range of saccade motor vectors. This ensures that a single visual input has access to a rich repertoire of motor outputs rather than being funneled toward a single one. This framework predicts that tasks that favor direct stimulus-to-saccade mapping could preferentially recruit regions of strong local RF-MF alignment, supporting rapid sensory-to-motor transformations. Conversely, tasks that require flexibility (such as anti-saccades or movements defined relative to a cue) could engage regions with divergent RF-MF pairings, with task rules biasing competition toward motor vectors appropriate to the instructed transformation. Thus, rather than implementing visuomotor flexibility solely through time-varying activity on a 1-to-1 map, the present findings indicate that a constrained mismatch in spatial scale between overlapping visual and motor mosaics generates a moiré topography that embeds the available repertoire of RF-MF pairings, providing a structural basis for flexible transformations between visual inputs and saccade outputs.

## Acknowledgements

Funding was provided by the Fiona and Sanjay Jha Chair in Neuroscience (J.H.R.) and NIH NEI grant R01EY028723 (J.H.R.)

## Methods

### Recording procedures

Two male marmosets (Monkey F and Monkey H) were implanted with a headpost, followed by a recording chamber. The recording chambers were implanted in the left and right hemispheres for Monkey F and Monkey H, respectively. The headpost and chamber implantation procedure has been previously published in Dotson et al. (2024). All experimental methods were approved by the Institutional Animal Care and Use Committee of The Salk Institute for Biological Studies. Marmosets are trained to enter custom-made marmoset chairs, which is then positioned in front of a calibrated and gamma-corrected LCD monitor (ASUS VG248QE; 100 Hz refresh rate). Eye position is measured using a video-based eye tracking system (ISCAN ELT-200, 500Hz sampling rate). MonkeyLogic is used to control the tasks and record behavioral data (Asaad & Eskandar, 2008; Hwang et al., 2019). Neuropixels recordings are acquired using the PXIe Acquisition Module (Neuropixels), and SpikeGLX acquisition software (for more details, see Dotson et al. (2024)). We use either the chamber or a stainless steel bone screw placed in the skull as the ground, and the built-in tip reference on the Neuropixels probe as the reference signal. Signals are re-referenced to a common average for the current source density and spike-field analyses. Spike sorting is performed using Kilosort 2.0 and Phy (Pachitariu et al., 2016). Artifact clusters are deleted and clusters that are clearly over split, based on comparing waveforms, are re-combined manually.

### Visual RF and saccade MF mapping task

Visual RFs are mapped using briefly flashed 2dva stimuli covering a 22 by 22dva area of the visual field. The stimulus is a filled yellow circle 2dva diameter. Stimuli are centered at points positioned on a grid (−10:2:10 horizontal, and -10:2:10 vertical) which covers a 22 by 22dva area when taking into account the stimulus size. Marmosets do not readily maintain fixation for long durations. Rather than ending the trial if they broke fixation, we allowed them to freely make saccades during the mapping procedure. To account for changes in eye position, we used the relative position of the stimulus to calculate the visual RFs, and data for a stimulus was only used if the eye position was stationary leading up to the stimulus presentation. We then utilized their natural saccadic behavior to map saccade MFs. We tried multiple stimulus durations (30 or 50ms) and inter-stimulus intervals (70, 170, and 250ms), but did not observe any differences with the location of RFs. For most sessions, on all or roughly half the trials, the stimulus was presented for 30ms with a 70ms inter-stimulus interval. On several sessions, blocks of trials were run requiring the monkey to maintain fixation. Due to the tendency of the marmosets to break fixation these blocks resulted in fewer useable stimulus presentations, and we saw no clear differences in the RF locations when comparing to the trials where the monkey was allowed to saccade. Subsequently, all trials were combined.

### Reconstruction methods

Recording locations are estimated using a 3D model of the skull and brain made in Blender (the Blender Foundation) and has been previously described in Dotson et al. (2024). Briefly, to reconstruct the location of the recording chamber with respect to the cortex, we used a CT scan taken after the headpost and chamber were implanted, and then aligned a 3D model of the chamber. We then registered a 3D model of the marmoset brain (Liu et al., 2021; marmosetbrainmapping.org) inside the skull model. After digitally removing the skull we could estimate the location of each cortical area with respect to the chamber. To estimate the location of the probe after each recording session, the microdrive is mounted on a pedestal and chamber identical to the ones used for recordings, and a picture is taken from below (Dotson et al., 2024). This provides the location of the probe with respect to the chamber. This information can be used to identify the location on the brain model, providing an estimate of the cortical area the probe likely penetrated (Supp. Fig. 1A).

The probe angle was estimated using the chamber model as a reference plane. Then we drew a small cylinder perpendicular to this plane and aligned it to the location on the brain model corresponding to the recording location. This provided us with a digital estimate of the probe path within the brain. Next, we estimated two angles, an anterior-posterior angle using a sagittal slice and a dorsal-ventral angle using a coronal slice. Angles were calculated by drawing a tangent line on the surface of the brain model and then measuring the angle between this tangent and the estimated probe trajectory. The angles were subsequently used to compute the horizontal displacement as function of depth along the probe.

### Visual RF and saccade MF vector angle quantification

RFs and MFs were detected using Spatial Information Analysis. Spatial information (SI) was calculated on 2D firing rate maps. Firing rate maps are 2D maps of the average firing rate at each stimulus or saccade endpoint location. To account for differences in eye position, the eye position during the stimulus presentation or the initial starting point of each saccade was subtracted. For each stimulus presentation, the data from 40ms to 90ms after the stimulus onset was used, so each stimulus presentation was 50ms. For saccades, the data from 0m to 50ms locked to the saccade onset was used. The firing rate at each location was then: # spikes / (# stimulus presentations or saccades * 50ms). SI in bits/spike for each neuron was computed using the following equation (Skaggs et al., 1996; Dotson & Yartsev, 2021):

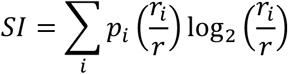

where *p*_i_ is the probability of the stimulus being in the i^th^ position, *r*_i_ is the firing rate of the neuron in the i^th^ position, and r is the average firing. Where r is computed as: *r* = ∑_*i*_ *p*_*i*_*r*_*i*_

A thorough discussion of the pros and cons of this method can be found in Souza et al. (2018). To assess the significance of the SI value for each neuron we compared the observed SI value to a shuffle distribution. The shuffle distribution was computed by randomly shuffling the spike responses with the stimulus or saccade locations. This maintained the same locations while randomizing the responses. We performed 1000 shuffles and tested whether the observed SI value was higher than 95^th^ percentile of those observed by chance (p<.05).

We identified the centroid of RFs and MFs using the built in Matlab (MathWorks) function regionprops. A 90% firing rate threshold (all pixels less than 90% of the max firing rate set to zero) was applied to firing rate maps prior to running regionprops. Only firing rate maps with a minimum of 2Hz firing rate were tested. Vector angles were calculated as the angle of the vector pointing to the centroid of each RF or MF. If more than one region was present, the region with the highest firing rate was used. Only penetrations with at least 10 RFs and 10 MFs were used for further analysis (18/39). These sites are within the FEF+ region (Supp. Fig. 1A).

To visualize spatial changes in RF and MF vector angles across cortex and to place all penetrations into a common analytical framework suitable for model comparison, we first smoothed the raw RF and MF angle measurements by locally averaging values within ±25 µm. This local averaging reduced high-frequency sampling noise while preserving mesoscale spatial structure. The smoothed angles were then linearly interpolated and resampled at uniform 50 µm intervals, producing regularly spaced angles that could be directly combined across penetrations and compared with model simulations. The distribution of angular differences between neighboring resampled points was computed and used as the primary statistic for fitting the mosaic models. This metric captures the rate and structure of angular change independent of absolute angle values. Because some neurons at the beginning or end of penetrations exhibited only RF or only MF tuning, the effective horizontal extents of the RF and MF maps occasionally differed slightly, resulting in small differences in line length after interpolation. To quantify RF–MF angular relationships, we computed the angular difference at each RF sample location relative to the nearest value from the interpolated MF profile. This nearest-neighbor matching ensured spatial alignment while avoiding extrapolation beyond measured regions.

### Mathematical mosaic map models

The annulus mosaic model (AM model) is a Fourier-based model in which the spatial pattern arises from the superposition of plane waves (Niebur & Wörgötter, 1994; Wolf & Geisel, 1998). The Fourier domain was represented as a two-dimensional grid of spatial frequency coordinates (*k*_*x*_, *k*_*y*_). The radial spatial frequency was defined as 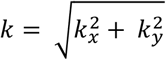, corresponding to the magnitude of the two-dimensional spatial frequency (wavevector). Mosaic maps were generated by specifying a set of Fourier components whose wavevectors were restricted to an annulus in 2D Fourier space, thereby constraining both orientation and spatial frequency content. A 2D array of random phase values *θ*(*k*_*x*_, *k*_*y*_), drawn uniformly from [0, 2*π*), was converted to complex unit-magnitude coefficients 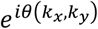 and multiplied pointwise by an amplitude mask *A*(*k*_*x*_, *k*_*y*_). The resulting complex Fourier spectrum was transformed back into the spatial domain using an inverse Fourier transform. The final mosaic map was defined as the wrapped phase angle of the resulting complex field after normalization by its magnitude, so that the resulting map depends only on phase. Formally, the mosaic map was defined as:

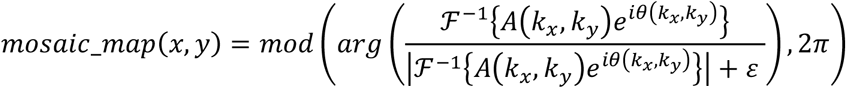

where ℱ^-1^{·} denotes the 2D inverse Fourier transform and *ε* is a small constant added to prevent division by zero. The amplitude mask *A*(*k*_*x*_, *k*_*y*_) was defined to select Fourier components within a narrow annulus in radial spatial frequency. Specifically, grid points whose radial spatial frequency satisfied *k*_0_(1 − *bw*) < *k* < *k*_0_(1 + *bw*), were assigned a value of 1, while all other grid points were assigned a value of 0. Here, *k*_0_ denotes the preferred spatial frequency index and *bw* the fractional bandwidth. Maps were generated on a square grid of 400 × 400 pixels. The grid was assigned a spatial extent of 4 × 4 mm, corresponding to a resolution of 10 µm per pixel. The preferred radial spatial frequency index *k*_0_ was varied between 1 and 20, corresponding to 0.25 to 5 cycles/mm for the 4-mm grid, and the fractional bandwidth *bw* was varied between 0.1 and 0.5. For each combination of *k*_0_ and *bw*, we made 1,000 independent mosaic maps by generating new random phases *θ*(*k*_*x*_, *k*_*y*_) for each.

The noise mosaic model (NM model) generated mosaic maps by convolving a complex Gaussian white-noise field with a 2D Gaussian kernel. A complex-valued white noise field was constructed by independently sampling the real and imaginary components from zero-mean Gaussian distributions. Spatial correlations were introduced by convolving this field with an isotropic Gaussian kernel with standard deviation σ (in pixels), truncated at three standard deviations and normalized to unit mass. The resulting complex field was then normalized to unit magnitude at each spatial location, preserving phase while removing amplitude variations. The orientation map was defined as the wrapped phase of the normalized field, yielding a spatially smooth, irregular mosaic. Maps were generated on a 400 × 400 pixel grid. Rather than varying a preferred radial spatial frequency or bandwidth (as in the AM model), spatial scale in the NM model was controlled exclusively by the Gaussian kernel width σ, which was systematically varied from 0.5 to 15 pixels corresponding to 5 to 150 µm (10 µm per pixel). For each value of σ, 1,000 independent realizations were generated by sampling new complex white-noise fields.

For both AM and NM models, analyses were performed in two stages. First, the visual and saccade mosaics were fit independently by comparing the distributions of within-map angular differences to those observed empirically. This procedure identified the narrow band of spatial scales that reproduced the locally smooth changes in angle punctuated by abrupt transitions characteristic of the data. Second, paired-map simulations were used to determine whether the structured RF-MF angle differences could emerge from mosaics with offset spatial scales and partial coupling. In the AM model, partial coupling arose from generating the visual and saccade maps using the same set of random Fourier coefficients while allowing their annular frequency bands to differ. When the annuli overlapped, shared Fourier components produced correlated structure between the maps. Thus, the degree of overlap in the center spatial frequency scaled the correlation between maps. To introduce controlled correlation between the visual and saccade NM maps, both maps were derived from a shared complex white-noise field, with additional independent noise added to one map to control the degree of coupling. For each permutation, a complex Gaussian field was sampled and used as the shared component for both maps. The second map additionally included an independent complex noise field. The shared and independent components were linearly mixed in proportions determined by a correlation parameter (ρ), which was systematically varied between 0 and 1. When ρ = 0 the maps were independent; when ρ = 1 they were fully shared prior to smoothing.

To compare model outputs to the empirical data, each simulated map (AM or NM) was sampled using randomly positioned and randomly oriented line segments (representing the horizontal sections of cortex that were sample by the probes) whose lengths matched those measured experimentally. Along each line, angle values were sampled at 50 µm spacing, and the absolute differences between neighboring samples were computed. The absolute angular differences were pooled across lines to generate a distribution of angle differences for each simulation. Model-generated distributions of angular differences were compared to the corresponding empirical distributions using two-sample Kolmogorov–Smirnov (K-S) tests. The K-S test evaluates the null hypothesis that two samples are drawn from the same continuous distribution. For each model realization, failure to reject the null hypothesis at α = 0.05 was recorded. For paired-map analyses, RF-MF difference maps were constructed by computing the absolute value of the angular difference between corresponding visual and saccade maps (interference maps), and the resulting distributions were compared to the empirical RF-MF difference distribution using the same procedure.

### Microstimulation methods

Marmosets were trained to fixate a small visual target (approximately 2.7 × 2.7dva) presented at one of five possible locations on the screen. The target was presented either at the center of the screen or at one of four peripheral locations corresponding to the corners of a square, with an eccentricity of 4.24dva. Each trial began with fixation on the target for 250ms, after which the target was extinguished. On approximately 50% of trials, electrical microstimulation was delivered at the time the fixation target was removed. In Monkey F, microstimulation was delivered following standard protocols using a single Pt/Ir microelectrode (Tehovnik et al., 2000). The electrode was inserted at a slight angle relative to the cortical surface and advanced in small, regular steps to stimulate multiple sites along each penetration. Stimulation was delivered using a Grass S88 stimulator coupled to two PSIU-6 stimulus isolation units. Microstimulation consisted of cathode-leading, charge-balanced biphasic current pulse trains delivered for 100ms at 200 Hz, with phase durations of 0.2ms and current amplitudes of 50 or 100 µA. In Monkey H, microstimulation was delivered through individual contacts on a 64-channel laminar probe (50 µm contact spacing; Atlas Neuro) coated with PEDOT:PSS to enhance stimulation efficacy. Stimulation was delivered using an RHS Stim/Recording System (Intan Technologies), which was used for both neural recording and stimulation. Microstimulation consisted of cathode-leading, charge-balanced biphasic current pulse trains delivered for approximately 50ms at 300 Hz, with phase durations of 0.2ms and a current amplitude of 25 µA. The location of recording sites and the calculation of the angle of the penetrations was performed in the same manner as the Neuropixels recordings.

## Supplemental Figures

**Supp. Fig. 1.**
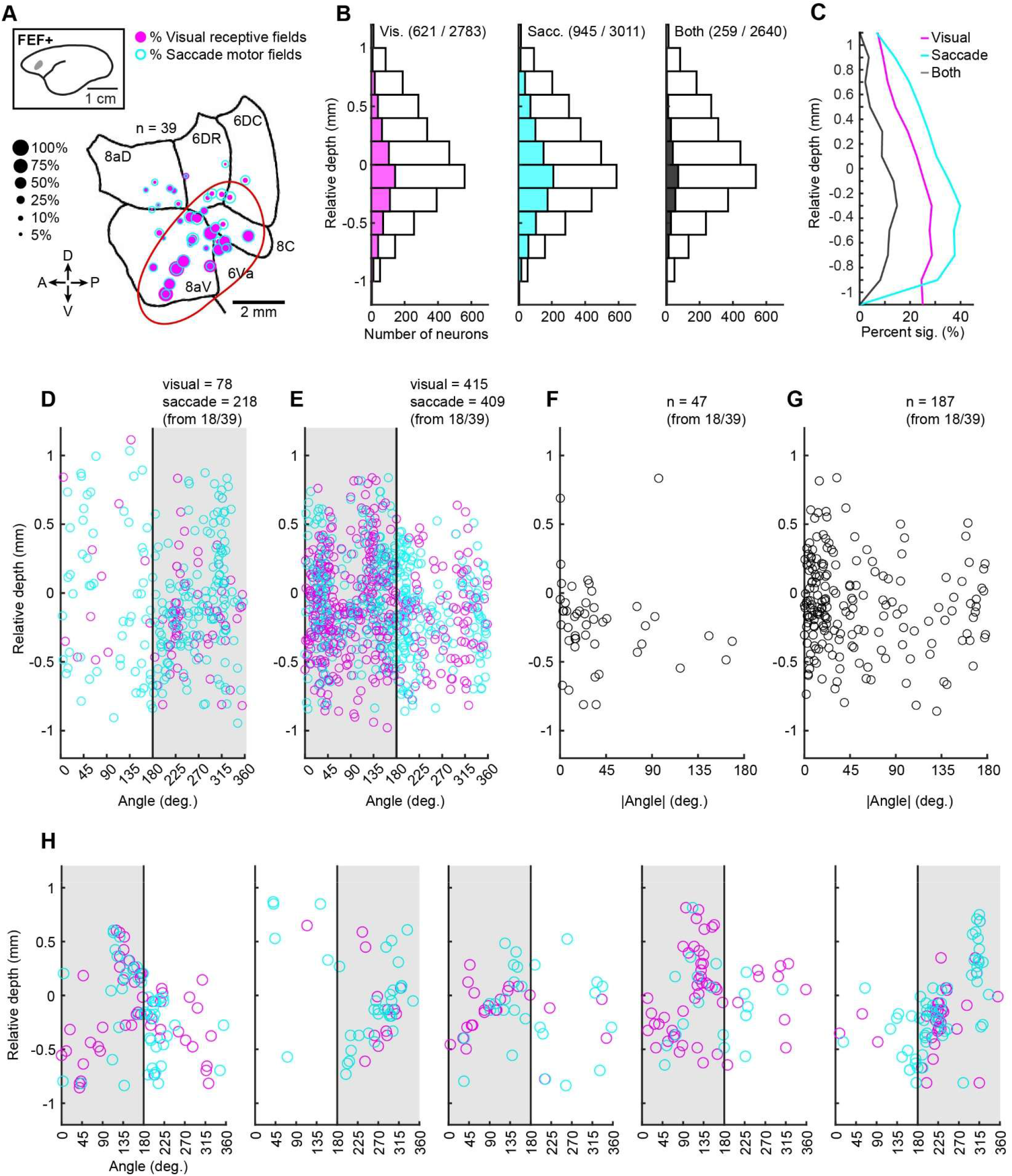
Estimated locations of recording sites and summary of RF and MF incidence and vector angles by relative cortical depth. **A**. Incidence of visual RFs (magenta) and saccade MFs (cyan) shown at the estimated locations of recording sites. Data is combined across both monkeys. Recordings in Monkey F and Monkey H were in the left and right hemispheres, respectively. The red oval indicates the estimated FEF+ region. **B**. Incidence of visual RFs, saccade MFs, and both as a function of depth. White bars indicate the total number of neurons. The inclusion criterion (>2Hz firing rate) is applied separately to each analysis; as a result, the total number of neurons differs across analyses. **C**. Percentage of neurons with RFs, MFs, and both as a function of depth (same data as B). **D, E**. Distribution of RF and MF angles by relative depth for Monkey F (D) and Monkey H (E). Angles are defined with 0° (360°) at the top and increase counterclockwise so that 180° is at the bottom. Under this convention, 0°-180° corresponds to the left hemifield, whereas 180°-360° corresponds to the right hemifield. Shaded region indicates the contralateral hemifield. **F, G**. Distribution of the absolute angle differences between RFs and MFs for neurons with both, distributed by relative depth for Monkey F (F) and Monkey H (G). Only data for recording sites with at least 10 RFs and 10 MFs (18/39) are included for plots D-G. **H**. Examples of the RF and MF angles by relative depth for single penetrations. The first three panels correspond to the examples in Figure 2 E-G. Panels 1, 3, and 4 are from Monkey H, and the remaining panels are from Monkey F. The shaded regions correspond to the contralateral hemisphere.

**Supp. Fig. 2.**
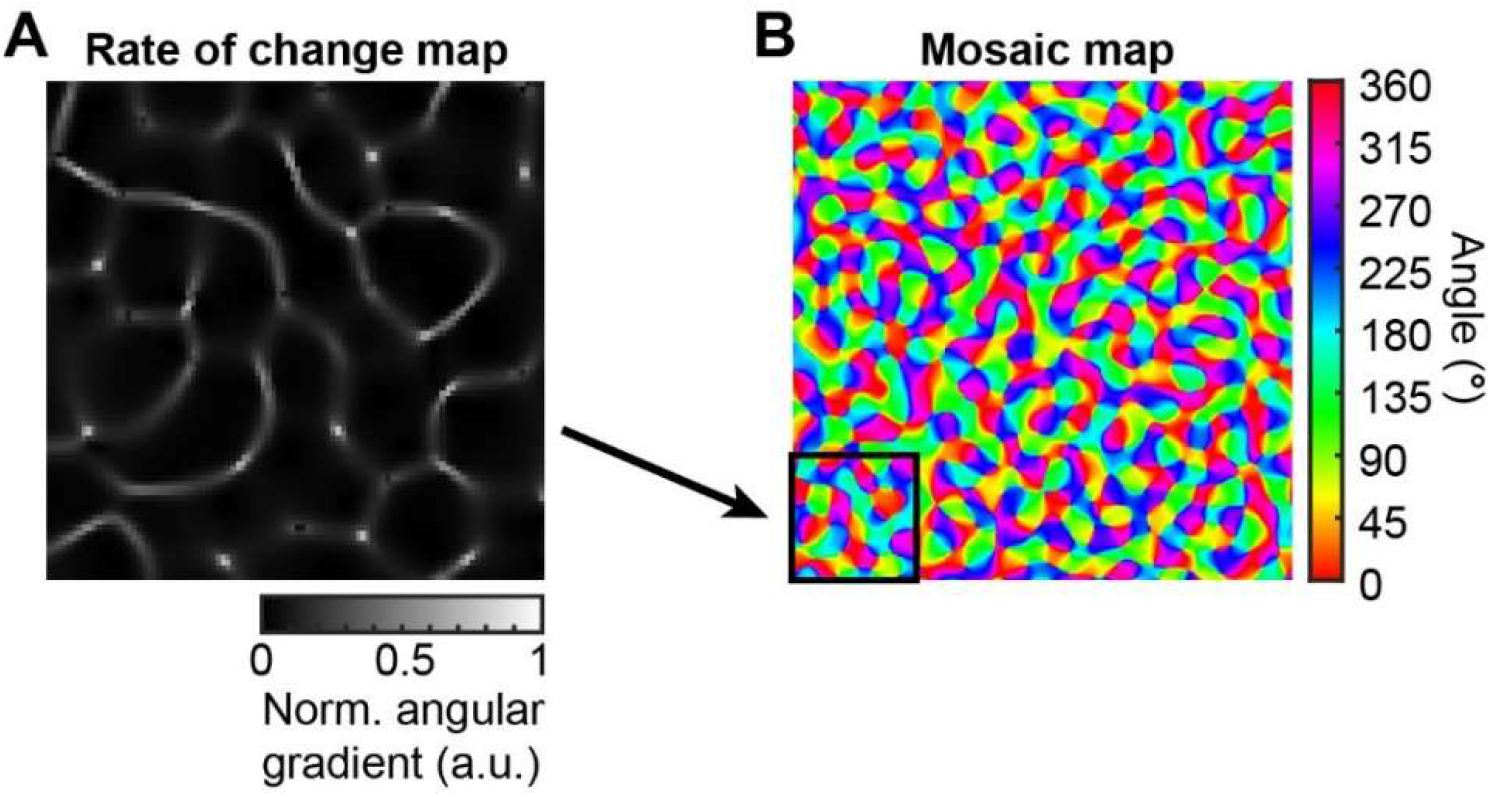
The AM model generates a biologically realistic rate of change map (i.e., Weliky et al., 1996). **A**. Normalized angular gradient computed from a portion of an example mosaic map (shown in **B**). The gradient magnitude quantifies the local rate of change in preferred angle across cortical space. Regions of low gradient reflect smooth, continuous variation in angular preference, whereas high-gradient regions (white curvilinear contours) mark abrupt transitions in angle, analogous to fracture lines.

**Supp. Fig. 3.**
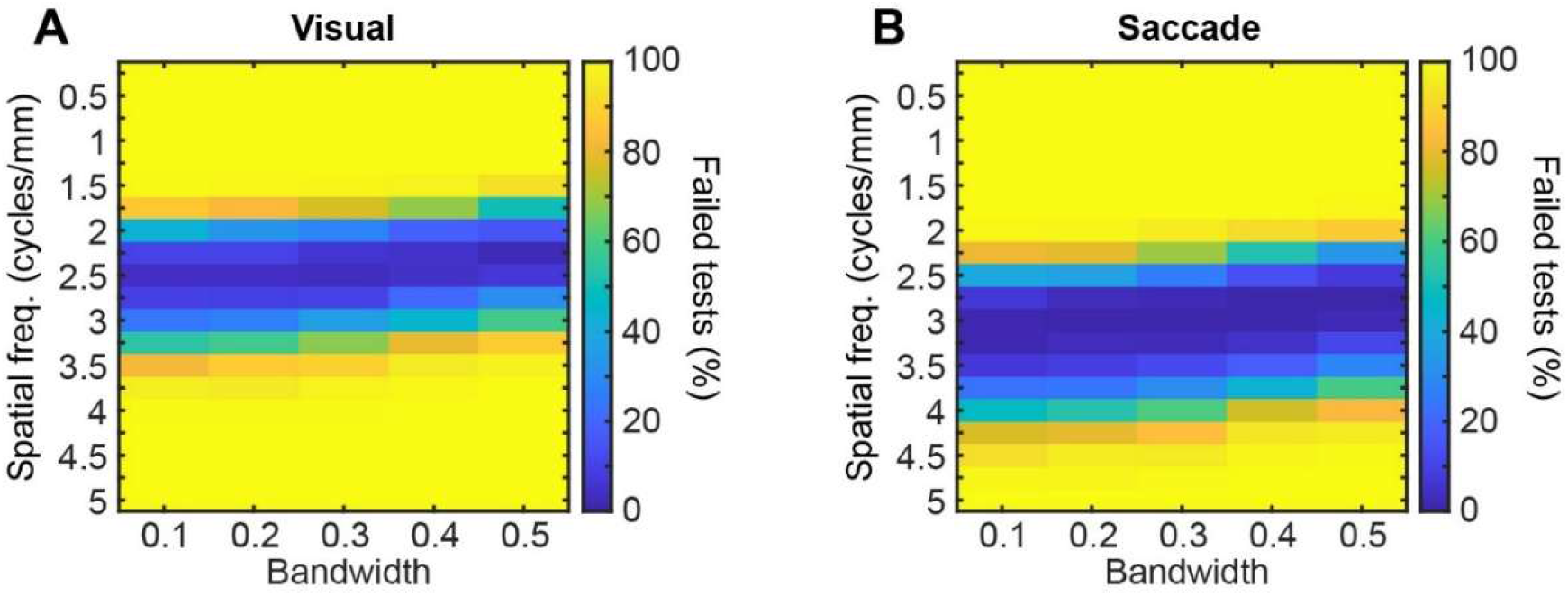
Bandwidth had a weak influence on the preferred spatial frequencies. We systematically varied the bandwidth parameter of the AM model to assess its influence on the inferred spatial scale of the mosaic organization. Across a broad range of bandwidth values, model fits were highly stable for both the visual (**A**) and saccade (**B**) maps. Although increasing bandwidth slightly reduced the preferred spatial frequency for both modalities, this effect was weak. Thus, the dominant factor governing the model fits is the central spatial frequency, with bandwidth exerting only a secondary influence. This robustness indicates that the inferred spatial structure of the visual and saccade maps reflects genuine features of the data rather than fine-tuned parameter choices.

**Supp. Fig. 4.**
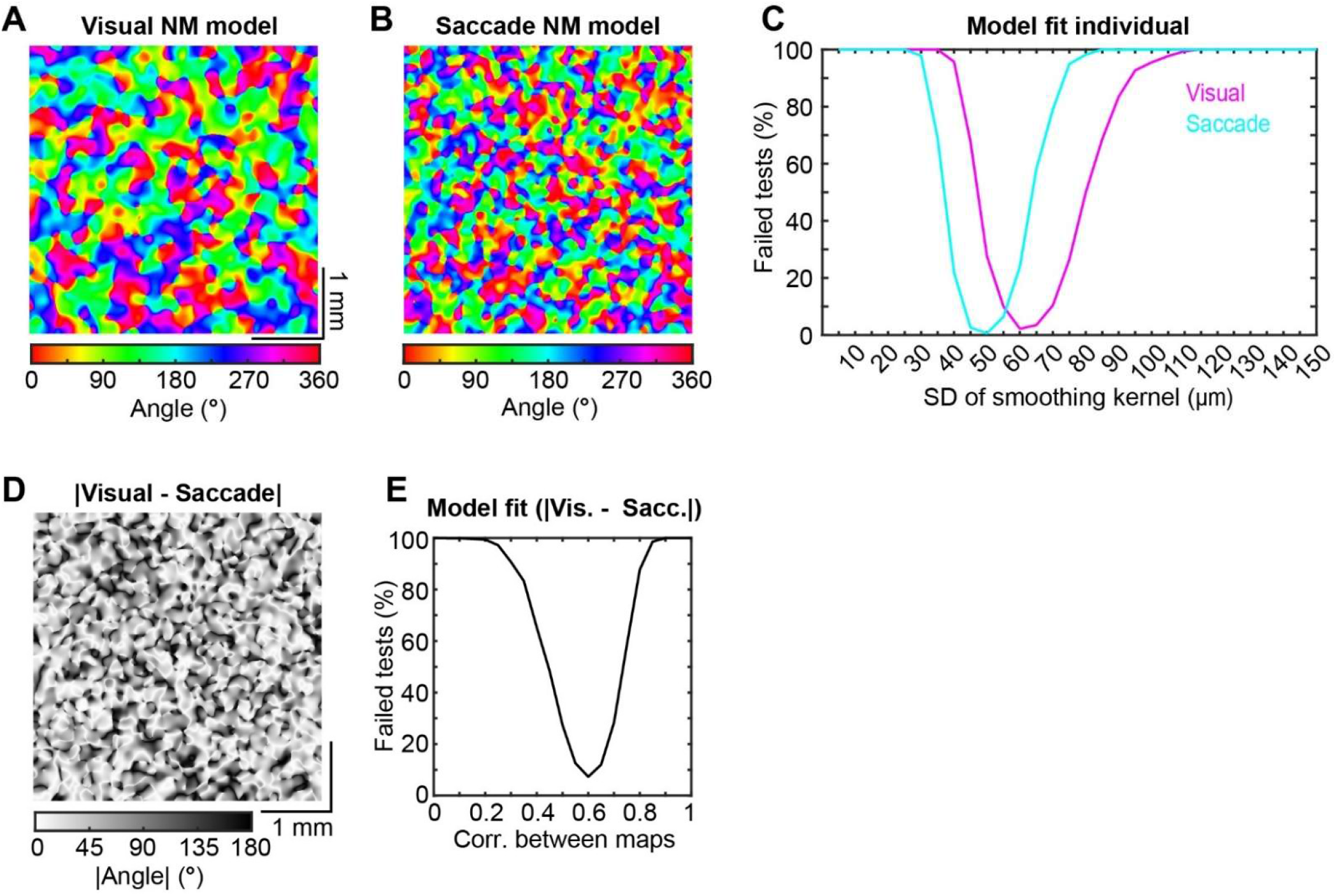
NM models replicates real data with partially correlated maps at different preferred spatial scales. **A, B**. Examples of randomly generated visual (SD = 50 µm) and saccade (60 SD = 60 µm) mosaic maps from the NM model. The example maps are generated with a correlation of 0.6. **C**. Results from testing the distribution of differences from the model to the real data at a range SD values. Only models within a narrow but offset range of SDs fit the data. **D**. Interference pattern obtained by subtracting the example visual and saccade maps (A, B). Despite the absence of periodic structure in the NM model, subtracting partially correlated maps with offset spatial scales produces a structured moiré-like pattern of alternating alignment and divergence. **E**. Model performance as a function of inter-map correlations for NM models with SD = 50 µm and SD = 60 µm. Uncorrelated and fully correlated maps failed to produce the differences between visual and saccade angles, whereas models with a correlation near 0.6 matched the data.

**Supp. Fig. 5.**
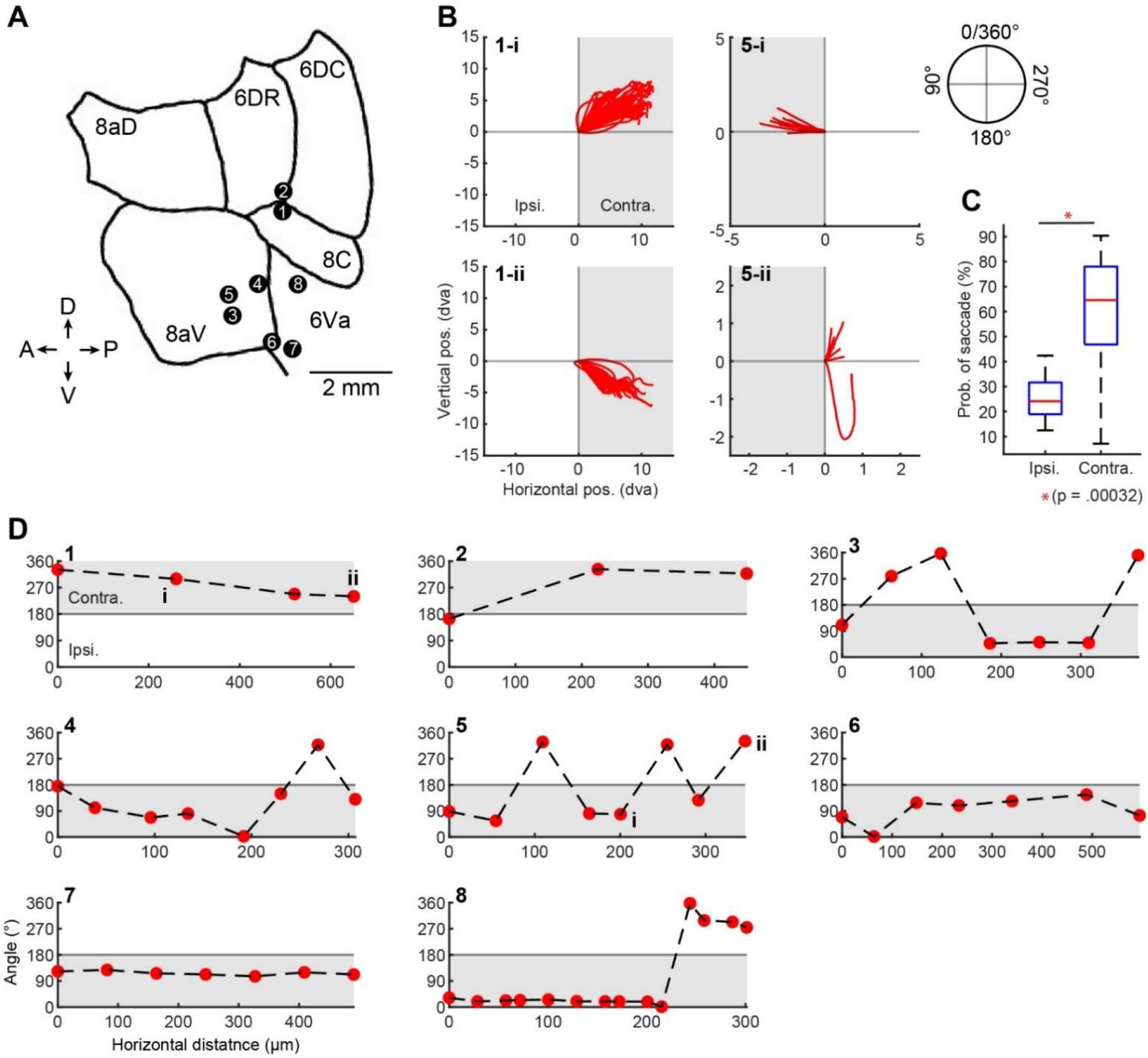
The angles of saccades driven by microstimulation are also consistent with a mosaic topography. We observe a similar pattern of smooth changes and abrupt jumps, consistent with a mosaic topography, and prior reports of microstimulation in macaques (Bruce et al., 1985). **A)** Estimated locations of penetration sites. Numbers indicate the associated plots in part D. **B)** Examples of saccades caused by microstimulation at different sites on two different recording sessions. The first column shows saccades being driven to the upper and lower contra-lateral hemifield. The second column shows saccades being driven to the contra and ipsilateral hemifields. The numbers match the plot number in part D, and the roman numerals indicate the site location. **C)** We found that the efficacy of saccade production was lower for the ipsilateral than the contralateral hemifield (Wilcoxon ranked sum test). **D)** Saccade angles for all microstimulation sessions. Red dots indicate the stimulation locations. Gray shading indicates the contralateral hemisphere. Plots 1 and 2 are from Monkey F where a single microelectrode was used, and the rest are from Monkey H where a 64-channel laminar probe was used.

**Supp. Fig. 6.**
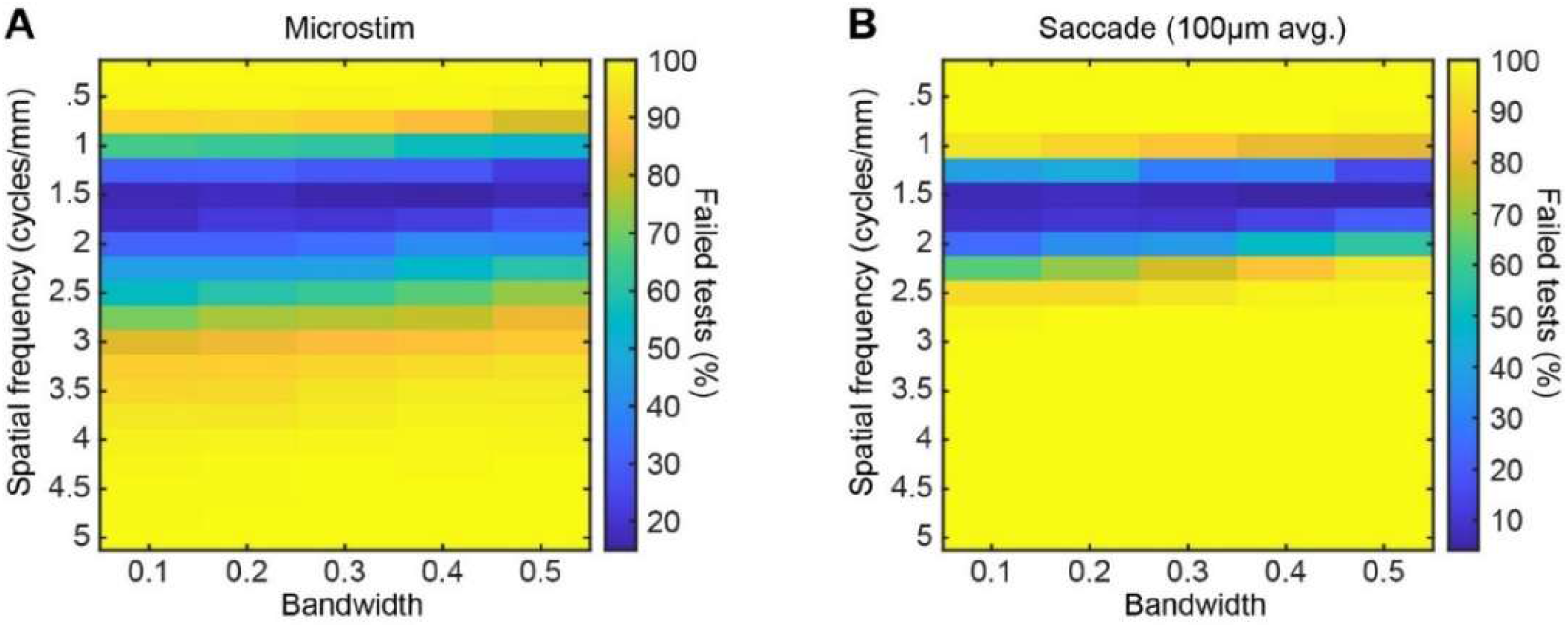
Results for mosaic map analysis on microstimulation data and saccade data with a larger spatial average. Using the same analysis applied to the RFs and MFs, we found that only a narrow range of spatial frequencies fit (A). However, the preferred SF range was lower than the preferred SF of the saccade MF angles. This could occur if the stimulation region is larger than the region used to measure the saccade angles, so we increased the region over which the saccade angles were averaged from 50 µm (±25 µm) to 100 µm (±50 µm). Due to the low spatial resolution of the stimulation sites for Monkey F, we only used data from Monkey H. After this modification, the preferred spatial frequency range of the saccade MF angles closely matched the microstimulation results (B).

